# Spatial and molecular insights into microglial roles in cerebellar aging

**DOI:** 10.1101/2025.03.01.640978

**Authors:** Andy P. Tsai, Douglas E. Henze, Eduardo Ramirez Lopez, James Haberberger, Chuanpeng Dong, Nannan Lu, Micaiah Atkins, Emma K. Costa, Amelia Farinas, Hamilton Se-Hwee Oh, Patricia Moran-Losada, Yann Le Guen, Alina Isakova, Stephen R. Quake, Tony Wyss-Coray

## Abstract

Aging induces region-specific functional decline across the brain. The cerebellum, critical for motor coordination and cognitive function, undergoes significant structural and functional changes with age. The molecular mechanisms driving cerebellar aging—particularly the role of cerebellar glia, including microglia—remain poorly understood. Here, we used single-nuclei RNA sequencing (snRNA-seq), microglial bulk RNA-seq, and multiplexed error-robust fluorescence *in situ* hybridization (MERFISH) to characterize transcriptional changes associated with cellular aging in the mouse cerebellum. We discovered that microglia exhibited the most pronounced age-related changes of all cell types and that their transcriptional signatures pointed to enhanced neuroprotective immune activation and reduced lipid-droplet accumulation compared to hippocampal microglia. Furthermore, cerebellar microglia in aged mice, compared to young mice, were found in closer proximity to granule cells. This relationship was characterized using the newly defined neuron-associated microglia score, which captures proximity-dependent transcriptional changes and suggests a novel microglial responsiveness. These findings underscore the unique adaptations of the cerebellum during aging and its potential resilience to Alzheimer’s disease (AD) related pathology, providing crucial insight into region-specific mechanisms that may shape disease susceptibility.

## Main

Aging is a primary driver of organ dysfunction^1^ and the leading risk factor for neurological disorders^2^. Understanding the mechanisms of brain aging is essential for developing strategies to combat age-related neurodegeneration, ultimately enhancing quality of life and extending cognitive health. This process is complex, as different brain regions age at varying rates, with some areas showing greater vulnerability to age-related changes^3,4^. The cerebellum, which contains approximately 70-80% of the brain’s neurons^5^ and is critical for motor coordination, sensory integration, and cognitive processing^6,7^, is among the regions most affected by aging, exhibiting structural and functional decline^3^. This deterioration contributes to an increased risk of falls, driven by impairments in balance, motor function, and fine motor skills.

Despite these vulnerabilities, the cerebellum demonstrates remarkable resilience to Alzheimer’s disease (AD) pathology^8,9^. Amyloid plaques are largely absent during early stages, and dystrophic neurites are rarely observed throughout the disease progression^10^. This resilience raises intriguing questions about the molecular and cellular mechanisms that confer protection against AD pathologies in the cerebellum. However, the lack of detailed studies on glial cell involvement in cerebellar aging has limited our understanding of age-related adaptations in this brain region, even though microglia display a high expression of AD risk genes^11,12^.

Cerebellar microglia are essential for maintaining neuronal homeostasis, with their high basal clearance activity likely mitigating cellular debris accumulation and preserving brain integrity^13^. This activity, distinct from other brain regions, may drive a unique microglia-mediated neuronal attrition, providing an adaptive mechanism to sustain cerebellar function during the natural decline in neuronal numbers observed in late adolescence^14^. Despite their importance, glial cell gene expression and spatial organization, including that of microglia, remain poorly characterized during healthy aging, leaving a significant gap in our understanding of how the cerebellum adapts to age-related changes. The cerebellum’s evolutionary conservation across vertebrates, coupled with its pronounced age-related changes, positions it as a unique model for studying brain aging^15^. However, it remains unclear whether molecular alterations in this highly conserved region can be leveraged to identify targets of brain aging. Addressing these gaps is critical for uncovering the mechanisms underlying cerebellar resilience to AD and understanding their broader role in age-related changes across the brain.

## Results

### Distinct transcriptional aging patterns in cerebellar cells

Cerebellar aging remains poorly understood, with limited insight into its molecular alterations. To address this, we isolated and analyzed the transcriptome of nuclei from the cerebellum of two female and two male mice each at 3, 12, 18, and 24 months of age. NeuN-negative and NeuN-positive nuclei were sorted to enrich for glial cell types, followed by single-nuclei RNA sequencing (snRNA-seq) of 90,164 individual nuclei. Uniform manifold approximation and projection (UMAP) analysis identified 15 transcriptionally distinct clusters, each representing a major cerebellar cell type (**Fig.1a**). These clusters were annotated based on the expression of well-established molecular markers specific to cerebellar cells^16^ (**Fig.1b**). To characterize cell type-specific transcriptional aging signatures, we performed differential expression analysis of snRNA-seq data, comparing the 3-month-old group to older age groups (12-, 18-, and 24-month-old). This analysis revealed distinct trajectories of differentially expressed genes (DEGs) for each cell type over time. Notably, microglia exhibited the most pronounced changes among the various cell types, particularly in the oldest cohort of 24-month-old mice, as shown by the average log fold change of the top ten upregulated DEGs (**Fig. 1c**). To further identify aging-associated genes, we applied the Mfuzz algorithm to cluster pseudo-bulk snRNA-seq data, which uncovered nine distinct gene expression trajectories across time points (**Fig. 1d**). Genes within clusters 1 and 7 demonstrated a strong correlation with aging, showing increasing expression levels with age (**Fig. 1e**). Remarkably, these age-associated genes were predominantly expressed in microglia (**Fig. 1f**), suggesting this cell type as the most transcriptionally responsive to aging. These findings highlight distinct, cell-type-specific transcriptional adaptations within the cerebellum during aging, with microglia emerging as prominent responders to age-related functional changes.

**Fig 1.**
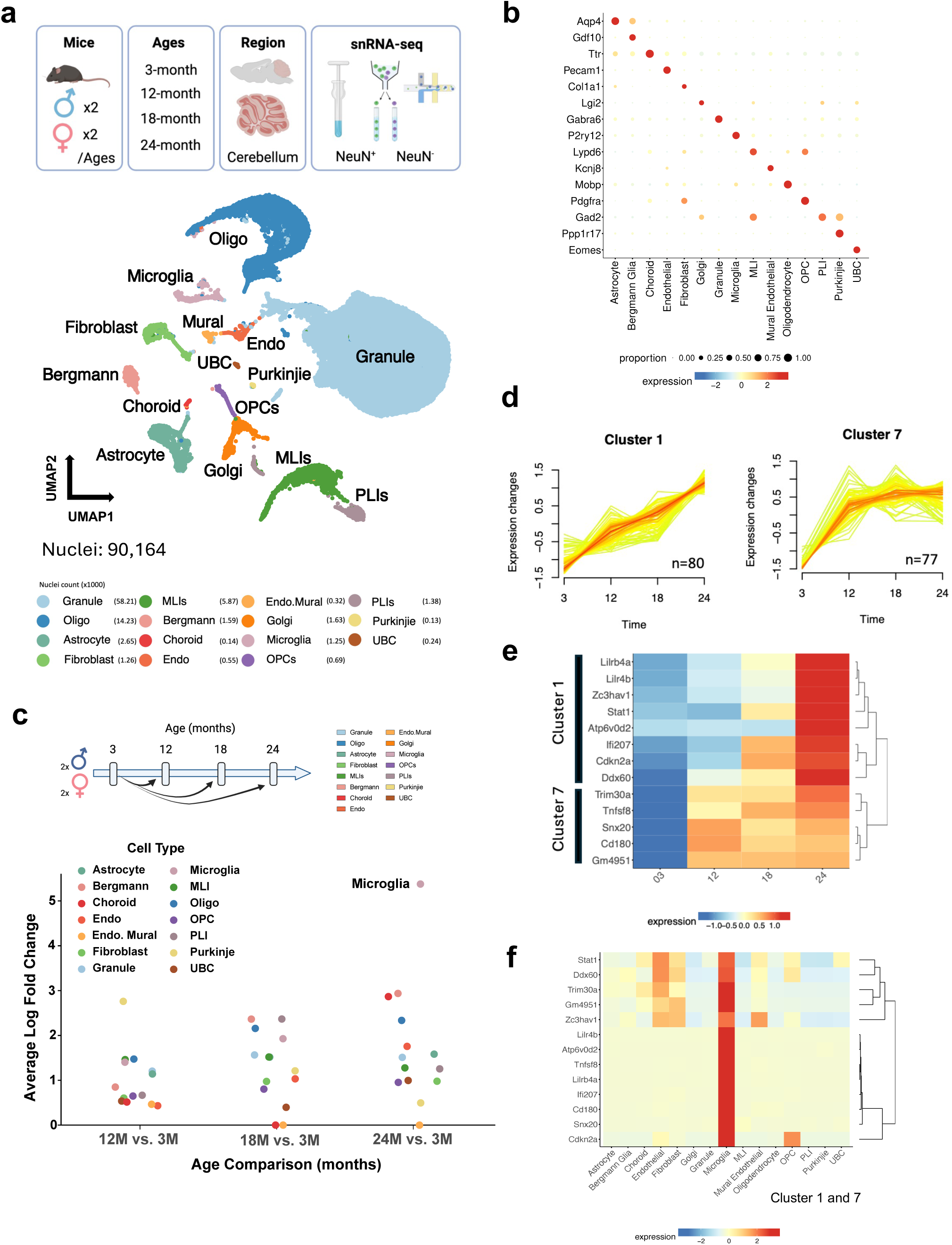
Distinct transcriptional aging patterns in cerebellar cells. **(a)**, Schematic representation of the experimental workflow for isolating and enriching glial nuclei from cerebellar tissue of 3-, 12-, 18-, and 24-month-old mice, followed by single-nuclei RNA sequencing (snRNA-seq). Uniform manifold approximation and projection (UMAP) visualization of 90,164 nuclei, colored by cell type, with corresponding counts listed. (**b)**, Dot plot of scaled expression levels for selected marker genes used to define cell types. (**c)**, Scatterplot illustrating the average fold change of the top ten differentially expressed genes (DEGs) for each cell type, with comparison made between each age group and 3-month-old mice. Dots are color-coded to represent cell types. (**d**) Age-dependent cerebellar gene expression changes are grouped into nine distinct clusters, with clusters 1 and 7 showing progressively increasing expression levels with age. (**e-f**) Heatmaps showing the expression levels of top genes from clusters 1 and 7 across four age groups (**e**) and among 15 cell types (**f**).

### Regional and age-associated transcriptional signatures of microglia

Acknowledging that different regions age at varying rates^3^, we investigate whether cerebellar microglia exhibit distinct transcriptional signatures compared to other brain regions. To address this, we examined the cortex (CTX), hippocampus (HP), thalamus (TH), hypothalamus (HTH), and striatum (STR). Microglia were isolated from these six regions of 1-, 3-, and 22-month-old male mice (n=6 per group), and transcriptomes were quantified using bulk RNA sequencing (RNA-seq) (**Fig. 2a**). Principal component analysis (PCA) revealed a major effect of age on gene expression across brain regions, with principal component 1 (PC1) scores increasing progressively with age. Notably, PC1 scores were highest in the cerebellum and lowest in the hippocampus and cortex, suggesting a unique transcriptional trajectory in cerebellar microglia during aging (**Fig. 2b-c**). Weighted gene co-expression network analysis (WGCNA) identified four key gene modules (Microglia M4, M9, M10, and M12) strongly associated with either age or brain region (**Fig. 2d**). Aging was linked to increased expression of the M4 and M12 modules and decreased expression of the M9 and M10 modules. These trends were most pronounced in cerebellar microglia, which exhibited the highest M4 module scores and the lowest M9 and M10 scores. Conversely, aged hippocampal microglia demonstrated higher M9 and M10 scores compared to their cerebellar counterparts (**Fig. 2e**). Gene Ontology (GO) enrichment analysis revealed the M4 module was enriched for type I interferon signaling pathway and cytokine-mediated signaling pathway, while the M12 module was linked to small GTPase mediated signal transduction. Receptor-mediated endocytosis was associated with the M9 module, and the M10 module was enriched for regulation of autophagy (**Fig. 2f)**. These biological processes are critically linked to the immune response, cellular signaling and homeostasis. For instance, type I interferon signaling (M4) plays a vital role in immune regulation, while autophagy (M10) maintains cellular integrity. In addition, small GTPase signaling (M12) and receptor-mediated endocytosis (M9) may influence cell communication and internalization, providing insights into molecular mechanisms of microglial aging and highlighting potential therapeutic targets for modulating these processes. To further identify microglial genes affected by age and region, we performed differential gene expression analysis on sorted microglia, comparing the cerebellum and hippocampus, as well as 22-month-old and 3-month-old groups. Genes that were differentially expressed in both comparisons including *Axl*, *Apoe*, *C4b*, *Lpl*, *Spp1*, *Hcar2*, and *Stat1* show distinct expression patterns across brain regions and age groups (**Fig. 2g**). To gain a deeper understanding of microglial functional states, RNA signatures from previous studies were used to calculate microglial scores (**Methods**). These included the neuroprotective activated plaque-responsive A and B (Act A & B) microglia score^17^ (**Fig. 2h**), lipid droplet (LD)-accumulating microglia score^18^ (**Fig. 2i**), senescence score and anti-senescence score^19,20^, disease-associated microglia score^21^, and activation score^22^ (**Extended Data Fig. 1a-d**). The Act A & B score, linked to amyloid plaque response and the neuroprotective effects of AD protective variant, was elevated in aged cerebellar microglia (**Fig. 2h**). In contrast, the LD-accumulating microglia score, associated with lipid metabolism, microglial dysfunction, and AD pathology, was significantly higher in the cortex and hippocampus (**Fig. 2i**). These findings suggest that regional differences in lipid droplet accumulation, a feature associated with *APOE4*-dependent neurotoxicity and neuroinflammation, may reflect distinct susceptibilities to aging and neurodegeneration.

**Fig 2.**
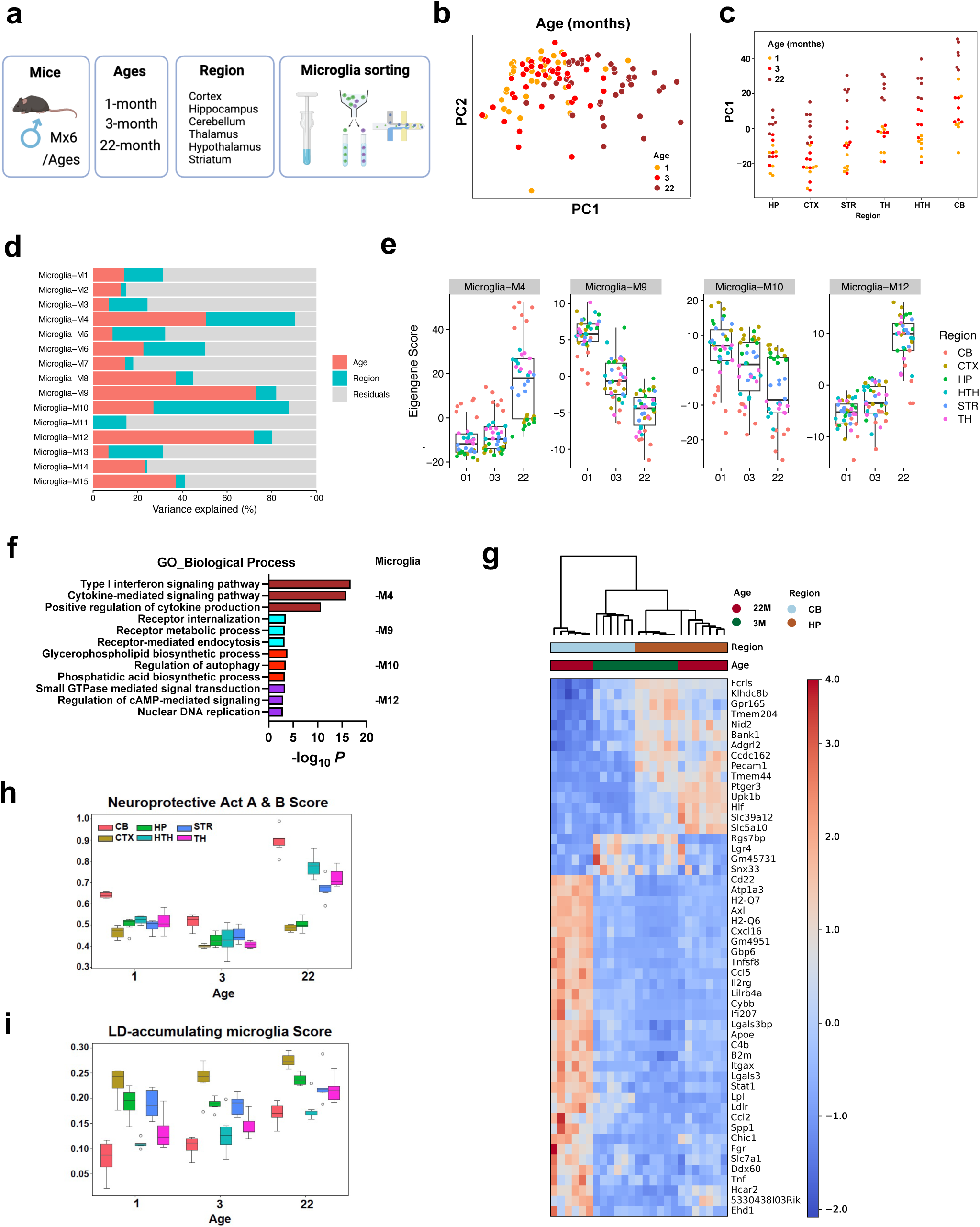
Regional and age-associated transcriptional signatures of microglia. **(a)**, Schematic of the experimental workflow for isolating microglia from six brain regions, cerebellum (CB), cortex (CTX), hippocampus (HP), hypothalamus (HTH), striatum (STR), and thalamus (TH), from 1-, 3-, and 22-month-old male mice. (**b)**, Principal component analysis (PCA) plot showing microglial transcriptional profiles differences across age groups and brain regions. (**c)**, Line plot of PC1 values across six brain regions, illustrating age-related transcriptional changes and unique aging trajectories of microglia. **(d)**, Bar chart depicting the percentage of variance explained by 15 microglial co-expression modules identified using weighted gene co-expression network analysis (WGCNA), highlighting the contributions of age and brain region. (**e)**, Box plots of eigengene scores for microglial modules M4, M9, M10, and M12, demonstrating region- and age-specific expression patterns across three age groups. (**f)**, Bar chart of top enriched Gene Ontology (GO) biological processes for genes within microglial modules M4, M9, M10, and M12. (**g)**, Heatmap showing top differentially expressed genes (DEGs) across two comparisons: 22-month-old versus 3-month-old mice and cerebellum versus hippocampus. **(h-i)**, Box plot showing the neuroprotective Act A & B score (**h**) and lipid droplet (LD)-accumulating microglia score (**i**), across three age groups and six brain regions.

### Aged cerebellum is characterized by increased microglial proximity to granule cells

To investigate the distinct transcriptional response of cerebellar microglia during aging, we estimated their spatial proximity to neighboring cells, including neurons, and compared these patterns to hippocampal microglia. Multiplexed error-robust fluorescent *in situ* hybridization (MERFISH) was applied to quantify spatial organizations and transcriptional signatures within the cellular architecture of the aging cerebellum and hippocampus. A parasagittal view of the mouse cerebellar and hippocampal layers is shown in **Fig. 3a**. MERFISH imaging was performed on sagittal sections of six mouse brains per sex, including both young (3-month-old) and aged (24-month-old) mice, using a panel of 500 genes selected for their relevance to aging, neuroinflammation, cellular stress, and cell-type specificity across brain regions (**Supplementary Table 1**). Cell types were annotated by integrating data with the Allen Brain Cell Atlas^23^. MERFISH images of the identified brain regions, including basal ganglia, cerebellum, cortex, hippocampus, hypothalamus, thalamus, and midbrain, pons, and medulla (MPM) region, are colored and shown in **Extended Data Fig. 2a**. The dataset contained 991,315 cells spanning 36 major cell classes across these 7 identified brain regions (**Extended Data Fig. 2b)**, with microglial distribution quantified (**Extended Data Fig. 2c)**. We analyzed spatial organization in the cerebellum and hippocampus of 24-month-old mice compared to 3-month-old mice to investigate the changes in cell proportions within microglial cellular neighborhoods, defined by a radial distance of 30 microns. Representative images of the cerebellum and hippocampus along with cell type labeling are shown in **Fig. 3b**. In the aged cerebellum, compared to the young, we observed increased granule cell-associated microglia, while decreased molecular layer interneurons (MLIs)-, Bergmann glia (Bergmann)-, and oligodendrocytes (Oligo)-associated microglia (**Fig. 3c**). In contrast, the aged hippocampus exhibited increased microglia and all other cells in the microglial neighborhoods (**Fig. 3d**). MERFISH images of cerebellum and hippocampus from both 3-month-old and 24-month-old mice revealed distinct spatial patterns of microglial organization during aging (**Fig. 3e)**. The significant increase in the number of microglia in proximity to granule cells distinguished the cerebellum from the hippocampus, where a generalized rise in the number of cells within microglial neighborhoods was observed, highlighting region-specific spatial and molecular adaptations of microglia during aging. To quantify the observations of increased granule cell-associated microglia, we analyzed MERFISH images and measured the percentage of microglia across different cerebellar layers at each age, including the white matter area, granule layer, and molecular layer. Our analysis revealed an increased percentage of microglia in the granule layer of aged mice (24-month-old) (**Fig. 3f**). To further validate these findings, we performed immunofluorescence staining and assessed relative numbers of microglia in the cerebellar granule layer. In 24-month-old aged mice we observed increased coverage of IBA1-positive microglia in the GABRA6-positive granule cell area of the cerebellum compared to 3-month-old young mice. (**Fig. 3g and Extended Data Fig. 2d**). To investigate the changes in transcriptomic signatures associated with spatial organization, we analyzed differentially expressed genes in microglia from the granule layer of aged mice compared to other cellular layers. Notably, genes including *Cbln1*, *Cdk2*, *Cxcl2*, *Slamf9*, and *Adcy1*, were uniquely upregulated in the granule layer of aged mice (**Fig. 3h**). The changes in microglial transcriptomic signatures suggest a functional shift, with microglia supporting neuronal integrity rather than responding to neuroinflammation in a specialized, region-specific manner. These findings highlight a unique and adaptive microglial response to aging in the cerebellum.

**Fig 3.**
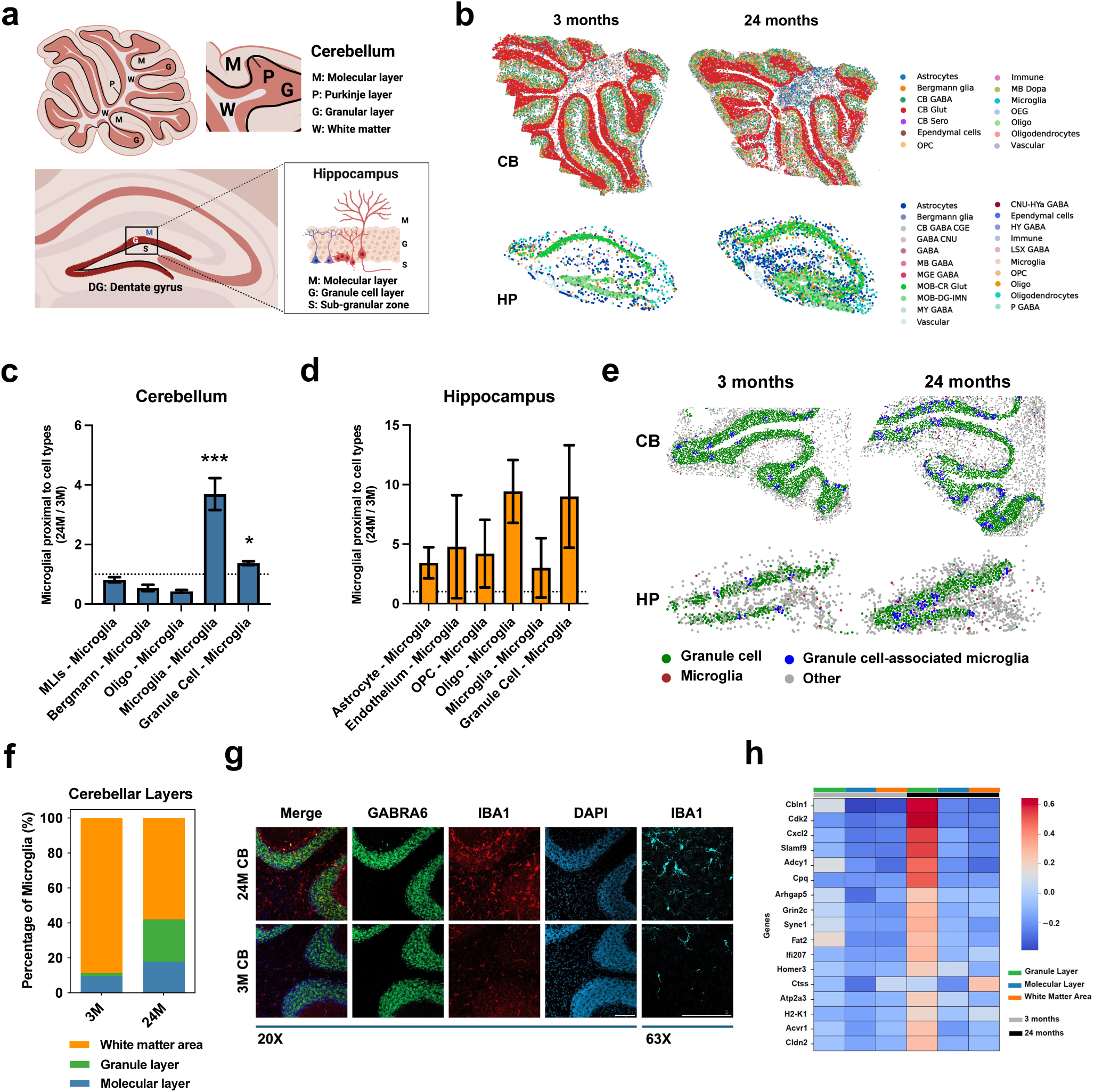
Aged cerebellum is characterized by increased microglial proximity to granule cells. **(a),** Schematic overview of the cellular layers of mouse cerebellum and hippocampus. **(b)**, MERFISH imaging of cerebellum and hippocampus from 3- and 24-month-old mice, colored by major cell types. **(c-d),** Bar plot showing the changes in microglial interactions with cerebellar (**c**) and hippocampal (**d**) cell types in aged versus young mice. **(e),** MERFISH imaging highlighting microglia (maroon), granule cells distant from microglia (green), granule cells near microglia (blue), and other cells (gray) of in the cerebellum and hippocampus of 3- and 24-month-old mice **(f),** Stacked bar plot showing the proportion of microglia within the molecular layer, granule layer, and white matter area of the cerebellum in 3- and 24-month-old mice. **(g),** Representative images of cerebellum from 3-month-old (3M) and 24-month-old (24M) mice, stained with GABRA6 to label granule cells (green), IBA1 to label microglia (red), and DAPI to label nuclei (blue). High-magnification IBA1 images (63x) are shown separately in cyan. Scale bar =100 μm. **(h),** Heatmap showing the top differentially expressed genes in microglia from the granule layer (green) compared to the molecular layer (blue) and white matter area (orange) in the aged cerebellum (black). All data are expressed as mean values ± SEM (*p<0.05, and ***p<0.001)

### Spatial proximity to granule cells shapes microglial aging in the cerebellum

Microglial signatures shaped by proximity to granule cells may indicate regional adaptations to brain aging. In young brains, microglia maintain a mosaic-like organization through balanced proliferation and apoptosis; however, aging disrupts this balance by altering cell cycle regulation and driving functional adaptations^24^. To investigate the effects of spatial organization on transcriptional changes, we scored the relative expression of functional gene sets in microglia, focusing on cell cycle-related genes and microglial genes associated with close proximity to granule cells (**Fig. 4a**). The cell cycle-related genes used to calculate the microglial cell cycle score regulate cell proliferation, DNA synthesis, and repair, and are typically activated in response to stress, injury, or debris clearance. We analyzed how the microglial cell cycle score (y-axis) is influenced by distance to granule cells (x-axis) in both the cerebellum (**Fig. 4b**) and hippocampus (**Fig. 4c**). Notably, the microglial cell cycle score showed a strong dependence on proximity to granule cells exclusively in the aged cerebellum, while no such dependence was observed in the hippocampus. Next, we identified genes with increased expression in microglia near cerebellar granule cells with MERFISH. We used these genes to calculate a newly defined “neuron-associated microglia score” in microglia. This score reflects a unique microglial response shaped by proximity to granule cells and their influence. Our analysis revealed that the influence of granule cells on microglia, as measured by the neuron-associated microglia score, varied across brain regions. This influence was most pronounced within 40 microns of granule cells, with the highest scores observed in aged cerebellar microglia (**Fig. 4d-e)**. We visualized the expression of genes contributing to the neuron-associated microglia score, including *Ctss*, *Slamf9*, *Cxcl2*, *Cdk2*, *H2-K1*, *Atp2a3*, and *Arhgap5*, in the cerebellum (**Extended Data Fig. 3a-m)** and hippocampus (**Extended Data Fig. 4a-m)**. The specificity of neuron-associated microglia score for microglia was evident in its gene expression patterns across individual glial cell types within 30 microns of granule cells in the aged cerebellum and hippocampus (**Extended Data Fig. 5a**). Supporting its validity, a significant increase in expression in the neuron-associated microglia score was observed in aged cerebellar microglia compared to young cerebellar microglia and other glial cell types in the aged cerebellum (**Extended Data Fig. 5b**). In addition, we applied the neuron-associated microglia score across five brain regions (**Extended Data Fig. 5c**), three cerebellar layers (**Extended Data Fig. 5d**) and sorted microglia bulk RNA-seq data from six regions across three ages (**Extended Data Fig. 5e**). These analyses demonstrated the utility of this novel neuron-associated microglia score in identifying regions with differential microglial responsiveness and capturing the effects of spatial organization on transcriptional changes in microglia during aging. Although the score was specifically identified in aged cerebellar microglia, the derived gene list serves as a versatile tool for understanding microglial function across diverse brain regions.

**Fig 4.**
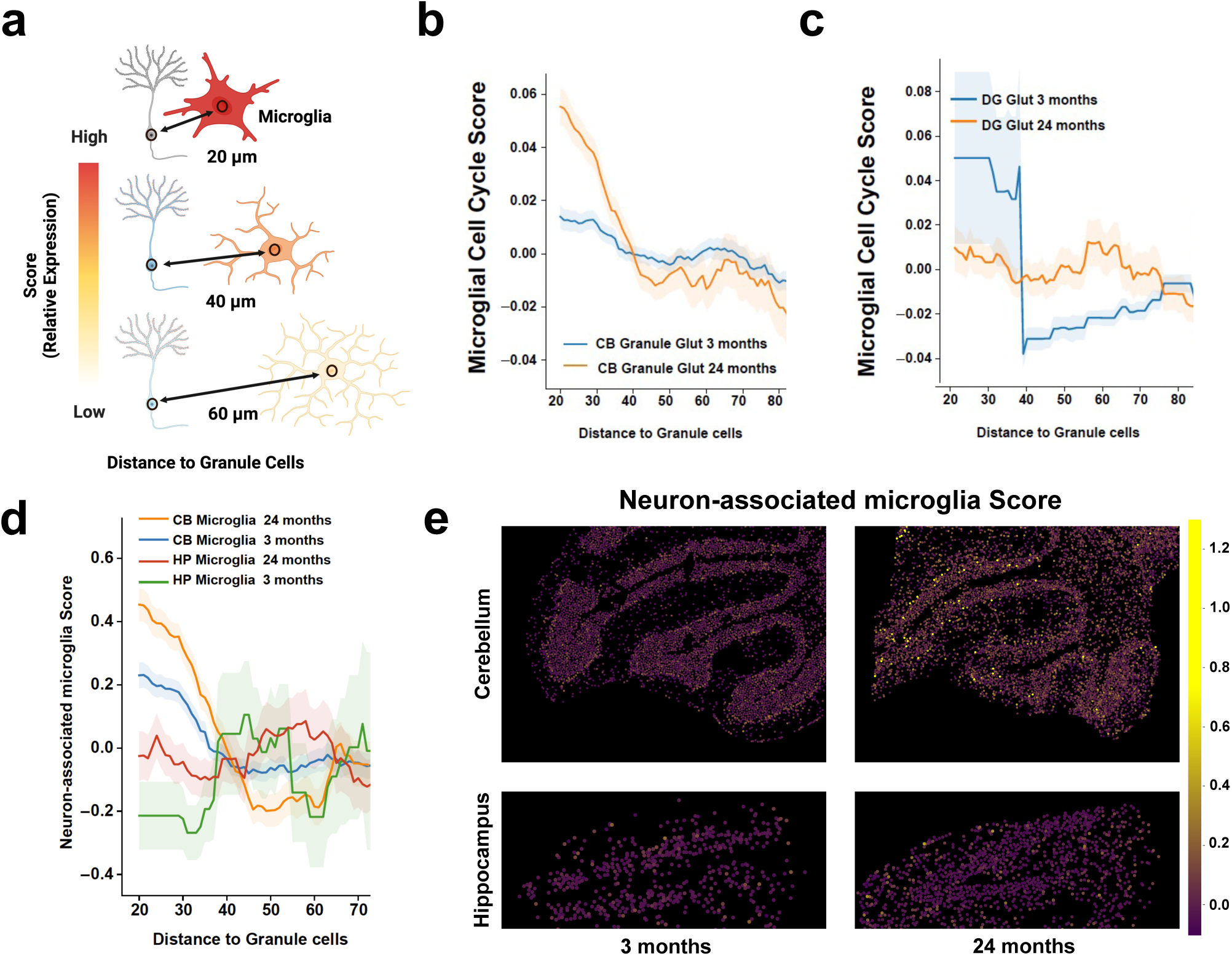
Spatial proximity to granule cells shapes microglial aging in the cerebellum. **(a),** Schematic illustrating the experimental workflow to examine the spatial distribution of microglial signatures and their proximity-dependent influence relative to granule cells. **(b-c),** Microglial cell cycle scores at varying distances from granule cells in the cerebellum **(b)** and hippocampal dentate gyrus **(c)** of 3-month (blue) or 24-month-old (orange) mice. **(d),** Line chart showing neuron-associated microglia scores at different distances from granule cells in the cerebellum and hippocampal dentate gyrus across age groups. **(e),** MERFISH images showing expression of neuron-associated microglia scores in the cerebellum and hippocampus of young (3 months) and aged (24 months) mice.

## Discussion

Age is the greatest known risk factor for AD, yet AD is not an inevitable consequence of biological aging. Despite extensive research attempting to explain the aging process^25^, the mechanisms that confer resilience to brain aging and related dementias remain poorly understood. Similar to distinct aging rates observed across organs^26^, brain regions show specific transcriptional changes with age, with the cerebellum being one the most significantly affected^3^. Given its strong transcriptional conservation across species^15,27^, the cerebellum offers a unique model for studying brain aging and the mechanism underlying age-related cognitive decline. However, detailed single-cell resolution data on glial cells and their age-related changes in the cerebellum were previously limited before this study.

Our study addresses these gaps by integrating cerebellar single-nuclei RNA-seq, sorted microglia RNA-seq, and MERFISH-based spatial transcriptomics to uncover transcriptional aging patterns in cerebellar cells. Among cell types in our dataset, microglia emerged as the most responsive cell type, showing region-specific adaptations that distinguish the cerebellum from other brain regions, particularly the hippocampus^28^. These findings provide new insights into region-specific aging mechanisms and highlight the potential resilience of the cerebellum to neurodegeneration.

Aging induced significant transcriptional changes in cerebellar microglia, characterized by upregulation of genes associated with immune activation, including *Axl*, *Lilrb4a*, *Cybb*, *Apoe*, *Lpl*, *Lgals3*, and *Hcar2*. These changes mirror the elevated microglial Act A & B score linked to AD protective variant^17,29^ (Fig. 2h) and disease-associated microglial signatures^21^ (Extended Data Fig. 1c). These transcriptional signatures could indicate that cerebellar microglia enhance the immune responses to facilitate debris clearance or aggregate removal, which may reduce neuronal stress and contribute to cerebellar resilience to neurodegeneration. In contrast, hippocampal microglia exhibited higher expression of lipid droplet (LD)-related genes, including *Acsl1*, *Nampt*, *Dpyd*, and *Cd163* (Fig 2i) ^18,30^, which have been implicated in *APOE4/4* genotype-dependent neuroinflammation and neurodegenerative vulnerability^18^. This divergence underscores region-specific adaptations to brain aging, with cerebellar microglia prioritizing immune activation, and hippocampal microglia maintaining a homeostatic state that may be less effective in mitigating age-related stress.

Spatial transcriptomics revealed a significant increase in microglial proximity to granule cells in aged cerebellum. distinguishing this region from the hippocampus and highlighting a unique spatial and molecular adaptation of microglia to aging. To quantify the effects of spatial organization on microglial function, we developed a neuron-associated microglia score, derived from microglial genes enriched near granule cells,. This score revealed strong proximity-dependent transcriptional changes in aged cerebellar microglia, which were absent in the hippocampus. This adaptation was associated with the upregulation of genes, including *Slamf9*, *Cxcl2*, and *Cdk2* in the granule layer, indicating potential specialized interactions aimed at supporting neuronal health and limiting neurotoxic inflammation. Although specialized and adapted to different demands, excitatory granule cells are present in both the cerebellum and hippocampus^16,31^, and have been implicated in cognitive impairments associated with aging^32,33^. The granule cell-rich architecture of the cerebellum likely facilitates focused interactions, enabling microglia to respond dynamically to age-related changes in their local environment. In addition, the upregulation of genes involved in disease-associated immune response^34^ and observed in microglia harboring the phospholipase C-gamma 2 variant (Neuroprotective Act A & B)^17^ further support the hypothesis that cerebellar microglia actively modulate granule cell health, contributing to neuroprotection. By contrast, hippocampal microglia adopt broader, less specific interactions with neighboring cells, reflecting a generalized aging response. This difference may help explain why the hippocampus is more vulnerable to neurodegeneration and underscores the importance of region-specific microglial adaptations in promoting brain resilience.

Cerebellar microglia adopt a distinctive approach to aging, characterized by region-specific immune activation and close proximity to granule cells. The elevated Act A & B score in cerebellar microglia emphasize their role in preserving neuronal integrity, as the associated genes have been previously linked to neuroprotective responses to amyloid plaques^17^. In addition, the lower LD accumulating microglia scores in the cerebellum compared to the hippocampus reflect a reduced susceptibility to lipid droplet accumulation, a major contributor to neurodegeneration in APOE4 carriers^18^. These region-specific differences highlight the enhanced ability of cerebellum to mitigate age-related stressors, and its resilience compared to other brain regions.

In conclusion, this study reveals a region-specific aging response in cerebellar microglia, highlighting the unique cellular and molecular strategies of the cerebellum in response to aging. These findings provide insight into potential protective mechanisms in the aging brain and suggest that targeted modulation of microglial function may support brain health and resilience to neurodegeneration in vulnerable brain regions.

## Methods

### Mice

C57BL/6J male and female mice (stock # 000664) were purchased from Jackson Laboratories. Up to five mice were housed per cage under specific-pathogen-free conditions in the Wu-Tsai Neurosciences Research Institute Veterinary Service Center at Stanford University School of Medicine. The colony room was maintained on a 12:12 h light/dark cycle with food and water provided ad libitum. Mice were euthanized by perfusion with ice-cold phosphate-buffered saline (PBS) following full anesthetization with Avertin® (125-250 mg/kg intraperitoneal injection). All animal care and experimental procedures complied with the Guide for the Care and Use of Laboratory Animals of the National Institutes of Health and were approved by Stanford University’s Administrative Panel on Laboratory Animal Care (APLAC).

### Nuclei Isolation and Fluorescence-Activated Cell Sorting (FACS)

Nuclei were isolated from the fresh-frozen cerebellar region of the left hemisphere of mice using Dounce homogenizers (Sigma-Aldrich, St. Louis, USA #D8938), and the Nuclei EZ Prep Kit (Sigma-Aldrich #Nuc101-1KT) according to previously published methods ^17^. The nuclei were then centrifuged at 300 g for 10 minutes at 4°C and resuspended in 50 μl of FACS buffer (1% BSA, 1× PBS; sterile filtered) containing 2 U/ml of Recombinant RNase Inhibitor (Takara #2313B) and anti-CD16/CD32 Fc block (BD Biosciences #553142). After incubation for 5-minutes, 50 μl of an antibody cocktail in FACS buffer containing 1 μl of NeuN antibody (Abcam #ab190565) and Recombinant RNase Inhibitor was added.

Samples were incubated with staining antibodies on ice for 30 minutes with gentle shaking, followed by centrifugation at 300 g for 10 minutes at 4°C. Nuclei were resuspended in 500 μl of FACS buffer containing Recombinant RNase Inhibitor and 1 μl of Hoechst 33342 was added. NeuN-positive and NeuN-negative nuclei were sorted separately into 1.5 ml DNA LoBind tubes containing 1 ml of final buffer (200 μl UltraPure BSA, Thermo #AM2618, 800 μl PBS, and 5 μl Protector RNase Inhibitor). A total of 50,000 single nuclei were sorted per sample. Nuclei concentrations were adjusted to ∼1 million nuclei/ml.

### Chromium 10X library generation and Illumina sequencing

Chromium 10X library generation and Illumina sequencing were conducted following previously published methods ^17^. Reagents for the Chromium Single Cell 3′ Library & Gel Bead Kit v3.1 (10X Genomics, Pleasanton, USA) were thawed and prepared following the manufacturer’s protocol. The nuclei/master mix solution was adjusted to target 10,000 nuclei per sample (5,000 from male mice and 5,000 from female mice) and loaded onto a standard Chromium Controller (10X Genomics) as instructed. All reactions, including library construction using the Chromium Single Cell 3′ Library Construction Kit v3, were performed according to the manufacturer’s protocol using the recommended reagents, consumables, and instruments. Quality control for cDNA and libraries was conducted using a Bioanalyzer (Agilent, Santa Clara, USA) at the Stanford Protein and Nucleic Acid Facility. Illumina sequencing of the 10X snRNA-seq libraries was performed by Novogene Co. Inc. (Sacramento, USA). Multiplexed libraries were sequenced using 2 × 150-bp paired-end reads in a single S4 lane on an Illumina NovaSeq S4 (Illumina, San Diego, USA), targeting 100 million reads per library. Base-calling, demultiplexing, and FASTQ file generation were performed for further data analysis.

### Single-nuclei RNA sequencing (snRNA-seq) analysis

The snRNA-seq reads were aligned to the mouse genome reference (mm10) using CellRanger (v7.1.0). Downstream analysis, including quality control (QC), cell clustering, and differential gene detection, was performed using the Seurat package. Specific QC criteria were applied as follows: 1) cells with less than 200 or more than 5000 detected features, or with more than 5% mitochondrial UMI counts, were removed; 2) integration was performed using LogNormalize on the top 3000 highly variable features, with default settings for other parameters; 3) clustering resolution was set to 0.6. UMAP was used for dimensionality reduction, and clusters were manually annotated based on canonical marker gene expression. These markers include *Gabra6* for granule cells (Granule), *Mog* and *Mobp* for oligodendrocytes (Oligo), *Aqp4* for astrocytes, *Col1a1* for fibroblasts, *Lypd6* for molecular layer interneurons (MLIs), *Gdf10* for Bergmann glia (Bergmann), *Ttr* for choroid plexus (Choroid), *Pecam1* for endothelial cells (Endo), *Kcnj8* for endothelial and mural cells (Endo.Mural), *Klhl1* and *Lgi2* for Golgi cells (Golgi), *Plcg2* and *P2ry12* for microglia, *Pdgfra* and *Olig2* for oligodendrocyte progenitor cells (OPCs), *Gad1* and *Gad2* for Purkinje layer interneurons (PLIs), *Ppp1r17* for Purkinje cells (Purkinje), and *Eomes* for unipolar brush cells (UBCs). Differentially expressed genes between cell types and conditions were identified using Seurat’s FindMarkers function.

Time-dependent genes were clustered using the R package Mfuzz, which applies soft clustering using the fuzzy c-means algorithm. The data of snRNA-seq were aggregated into pseudo-bulk values as input for Mfuzz clustering. The number of clusters was set to 9, and the optimal fuzzifier coefficient (m) was determined using the ‘mestimate’ function.

### Microglia isolation and bulk RNA-seq analysis

Microglia were isolated using ice-cold dounce homogenization, based on published methods ^35^. After perfusing mice with PBS, six brain regions (hypothalamus, thalamus, hippocampus, striatum, cortex, cerebellum) were dissected, chopped, and homogenized in cold medium A (HBSS, 15 mM HEPES, 0.5% glucose, and 1:500 DNase I). The homogenate was filtered and centrifuged. Myelin was removed with MACS myelin removal beads (Miltenyi) and MACS buffer (1x PBS, 2 mM EDTA, 0.5% BSA), followed by cell resuspension in FACS buffer (1x PBS, 2 mM EDTA, 1% FBS, 25 mM HEPES) with FC blocking (CD16/CD32). Cells were stained with CD11b-FITC, CD45-BUV395, and CD206-PE-Cy7, then washed, centrifuged, and dead cells excluded with Sytox Blue (1:1000). 6,000 microglia were sorted into RLT lysis buffer (Qiagen) with 1% 2-mercaptoethanol (Sigma, M6250) and frozen at −80 °C.

RNA was extracted with the RNeasy Plus Micro kit (Qiagen, 74034). cDNA and libraries were synthesized in-house using a modified Smart-seq2 protocol as previously described ^3,36,37^. Amplified cDNA was prepared for library sequencing, including bead clean-up, tagmentation, and indexing. Libraries were purified and assessed for quality using a Bioanalyzer (Agilent), then sequenced with Nextseq 550 high-output kit systems (Illumina, 20024907, paired end, 2 x 75 bp depth).

Principal Component Analysis (PCA) was conducted using scikit-learn package in python using normalized bulk RNA-seq data. Weighted Gene Co-expression Network Analysis (WGCNA) was used to identify gene modules correlated with traits of interest from bulk RNA-seq data. Data were normalized and filtered to retain highly variable genes. A soft-thresholding power was selected to create a scale-free network, and a topological overlap matrix (TOM) was used to cluster genes, identifying co-expression modules. Module eigengenes, representing module expression profiles, were correlated with traits to find significant associations. Gene Ontology (GO) enrichment analysis was performed on significant modules, and hub genes were identified based on module connectivity.

Gene set scores were calculated from microglial bulk RNA-seq data by averaging the expression of selected genes associated with the biological process of interest. Normalized expression values were used to calculate an average score per sample. Box plots were used to compare scores across conditions. Data processing and visualizations were performed in R using ggplot2 for boxplots. Neuroprotective Act A&B score ^17^ were calculated by genes including *Axl*, *Cd9*, *Csf1r*, *Hif1a*, *Itgax*, *Tmem163*, *Apoe*, *Cybb*, *Lilr4b*, *Lgals3*; LD-accumulating microglia score ^18,30^ were calculated by genes including *Slc25a5*, *Npl*, *Angptl7*, *Pde2a*, *Ldhb*, *Cd63*, *Sepp1*, *Sdcbp*, *Adipor1*, *Rbbp4*, *Cndp2*, *Hsd17b4*, *Hsd17b4*, *Gpd11*, *Dazap2*, *Hnmpk*, *Rapsn*, *Cat*, *Kl*, *Nampt*, *Acsl1*, *Dpyd*, *Cd163*; senescence score were calculated by genes including *Ccl2*, *Tgfb1*, *Il1b*, *Mmp3*, *Ccl5*, *Cxcl10*, *Serpine1*, *Cdkn2a*, *Glb1*, *Tnf*, *Il6*, *Cdkn1a*; Anti-senescence score were calculated by genes including *Cdk4*, *Rb1*, *Cdk2*, *Lmnb1*, *Mki67*, *Cdk6*; DAM score ^21,38^ were calculated by genes including *Trem2*, *Apoe, Tyrobp*, *Itgax, Clec7a*, *Lpl*, *Cst7*, *Spp1*, *Axl, Cd9*; Activation score ^22^ were calculated by genes including *B2m*, *Trem2*, *Ccl2*, *Apoe*, *Axl*, *Itgax*, *Cd9*, *C1qc*, *Lyz2*, *Ctss*, and Granule influence score were calculated by genes including *Cdk2*, *Cxcl2*, *Slamf9*, *Arhgap5*, *Ctss*, *Atp2a3*, *H2-K1*.

### MERFISH analysis

The annotated and pre-processed MERFISH data were obtained from Henze et al. ^39^. These data were used to analyze cell-cell interaction within the different brain regions at 3 and 24-month-old mice. As described previously ^40^, cells were only analyzed if they had a subclass label-transfer confidence score greater than 0.8. For each cell type pair within each region at each age, we then determined the number of each cell type contacting them and compared the difference between young and old contact pairs against the difference in the null distributions for each age. The null distributions were generated by locally shifting the positions of the cell types to random locations and calculating the cell-cell contacts amongst them. To consider two cells in proximity to one another, we established a distance threshold of 30 microns. We generated the null distribution by randomly shifting local cell positions within each brain region and counting cell-cell proximal pairs over 1000 rounds of permutations. To determine differences in cell-cell contacts, we fitted the distribution of differences between data from 3- and 24-month-old mouse samples contacts to the normal distribution and calculated the p-value for the ratiometric change between the ages to determine differences in the probability of cell-cell interaction with age.

To then analyze the number of microglia within each region of the cerebellum we analyzed only the cells of the cerebellum and for microglia that were closer to a Purkinje cell than to a granule cell we classified them as being from the molecular layer. The remaining cells were then divided amongst being in the granule layer versus in the white matter if the microglia were within 30 microns of a granule neuron, and the remainder of the microglia were delineated as white matter originating.

We then computed scores from the normalized and log-transformed gene expression values using the score_genes function in Scanpy. The genes used to calculate the cell cycle score include: ‘*Cdk4*’,’*Rb1*’,’*Cdk2*’,’*Lmnb1*’,’*Mki67*’, and ‘*Cdk6*’. The genes used to calculate the granule cell influence include: ‘*Cdk2*’, ‘*Cxcl2*’, ‘*Slamf9*’, ‘*Arhgap5*’, ‘*Ctss*’, ‘*Atp2a3*’, and ‘*H2-K1*’. The granule cell influence scores were selected from the differential expression between granule cell proximal microglia in the old compared to the young mice. We then computed these scores as a function of distance to granule cells either in the cerebellum (CB Granule Glut) or in the hippocampus (DG Glut) in both young (3 months) and old (24 months) mice. Each granule cell identified within 100 microns distance of a microglial cell was identified using kD-tree search in scikit-learn as explained previously ^22^. The average cell cycle or granule influence score was then calculated at one-micron steps from zero to seventy microns with a sliding twenty-micron window. The mean activation was also subtracted from these ranges so that they are centered around zero.

After establishing a granule cell influence score, we then performed differential gene expression analysis between young and old microglia that were within thirty microns of granule cells in either the cerebellum or the hippocampus. The differential gene expression between the cells was estimated using the Mann-Whitney-Wilcoxon test and sorted by log fold change from young to old.

### Immunofluorescence staining and image analysis

Perfused brains from 3- or 24-month-old mice were perfused and fixed in 4% paraformaldehyde for 24 hours at 4°C. After fixation, brains were cryoprotected in 30% sucrose at 4°C until fully infiltrated and then embedded. Brains were cut into 40-μm free-floating sections using a microtome. At least three paired brain sections per experiment were used for immunostaining. The sections were first washed and permeabilized in 0.1% Triton X-100 in PBS (PBST), then blocked with 5% normal donkey serum in PBST for 1 hour at room temperature (RT). Sections were incubated overnight at 4°C with primary antibodies in 5% normal donkey serum in PBST: Iba1 (goat, Novus Biologicals #NB100-1028, 1:1000), GABRA6 (rabbit, Synaptic Systems #224603, 1:500; AB_2564653), and Hoechst 33342 (Miltenyi Biotec #130-111-569, 1:10,000). After primary antibody incubation, sections were washed and incubated with appropriate species-specific AlexaFluor secondary antibodies (diluted 1:1000 in 5% normal donkey serum in PBST) for 1 hour at RT. Sections were then washed three times in PBST for a total of 30 minutes, counterstained, and mounted on slides. Immunofluorescence staining was based on published methods and images were captured using a fluorescence microscope with consistent exposure and gain settings across stains and animals ^41^.

Images were captured on a Leica Stellaris8 confocal microscope at high magnification and analyzed using ImageJ software (NIH); results were obtained from an average of at least three sections per mouse, with a threshold applied to all images. The threshold function in ImageJ was used to determine the percentage of immunoreactive area of the microglial marker (IBA1) within the granule layer (GABRA6), the region of interest (ROI). Plots were generated using GraphPad Prism (version 10.4.0).

## Data availability

Sequencing data are available through Gene Expression Omnibus under accession code: GSE290806.

## Code availability

Code used to analyze the datasets is available on GitHub at (https://github.com/doughenze/Aging_CB)

## Acknowledgments

The authors thank the past and present members of the Wyss-Coray lab, Quake lab, Chan Zuckerberg Biohub, and Knight Initiative lab for their feedback, valuable discussions, and support throughout the study. The authors also acknowledge all the animals sacrificed for this study. This work was supported by Larry L. Hillblom Foundation (A.P.T), the Milky Way Research Foundation (T.W.C), Simons Foundation (T.W.C), D. H. Chen Foundation (T.W.C), Knight Initiative for Brain Resilience (T.W.C), NIA R01 grant AG072255 (T.W.C), and Chan Zuckerberg Biohub (S.R.Q).

## Author information

These authors contributed equally: Andy P. Tsai, Douglas Henze.

Department of Neurology and Neurological Sciences, Stanford University School of Medicine, Stanford, CA, USA. (A.P.T); (E.R.L); (N.L); (M.A); (A.F); (H.S.O); (E.K.C); (P.M.L); (T.W.C). Wu Tsai Neurosciences Institute, Stanford University, Stanford, CA, USA. (A.P.T); (E.R.L); (N.L); (M.A); (A.F); (H.S.O); (E.K.C); (P.M.L); (A.I); (T.W.C). Department of Bioengineering, Stanford University, Stanford, CA, USA. (D.E.H); (S.R.Q). The Phil and Penny Knight Initiative for Brain Resilience, Stanford University, Stanford, CA, USA. (J.H); (A.I). Department of Genetics, Yale University School of Medicine, New Haven, CT, USA. (C.D). State University of New York Downstate Health Science University, College of Medicine, Brooklyn, NY, USA. (M.A). Quantitative Sciences Unit, Department of Medicine, Stanford University, Stanford, CA, USA. (Y.L.G). Department of Applied Physics, Stanford University, Stanford, CA, USA. (S.R.Q). Chan Zuckerberg Initiative, Redwood City, CA, USA. (S.R.Q).

## Author’s contribution

A.P.T, D.E.H, S.Q.R and T.W.C conceptualized and designed experiments. A.P.T, D.E.H, E.R.L, N.L, and M.A conducted experiments and collected data. A.P.T, D.E.H, E.R.L, J.H, C.D analyzed data. A.P.T, D.E.H, E.R.L, J.H, C.D, E.K.C, A.F, H.S.O, P.M.L, Y.L.G, A.I, S.Q.R and T.W.C discussed and interpreted data. A.P.T, D.E.H, E.R.L, J.H, C.D designed figures. A.P.T and T.W.C wrote the manuscript. A.P.T, D.E.H, S.Q.R and T.W.C supervised, directed, and managed the study. All authors discussed the results and commented on the manuscript.

## Competing interests

The authors declare no competing interests.

## Extended Data Figures

**Extended Data Fig 1.**
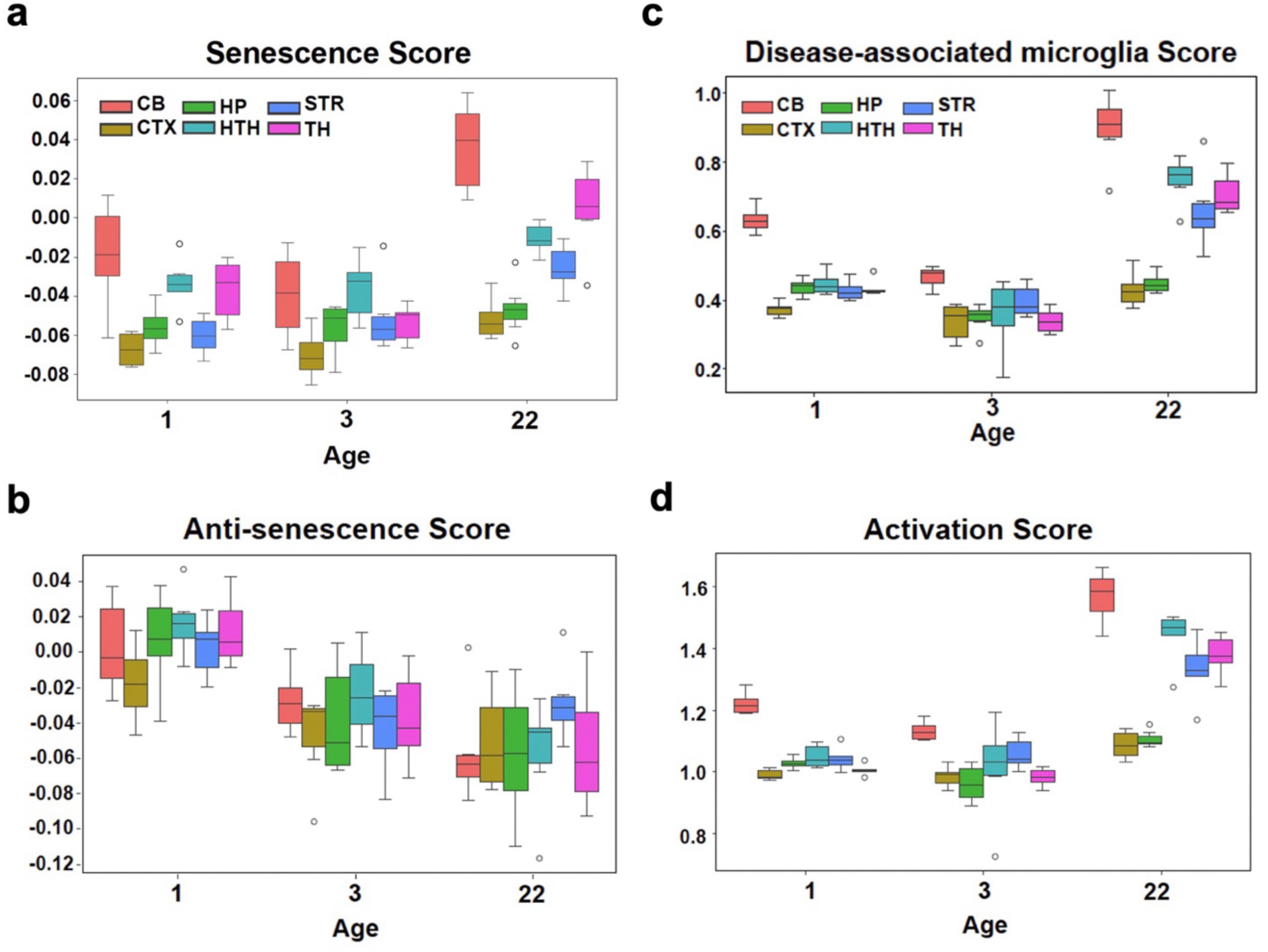
Microglial scores across ages in six brain regions. **(a-d),** Box plots showing senescence score **(a)**, disease-associated microglial score **(b)**, anti-senescence score **(c)**, and activation score **(d)** of microglia across three different ages and six brain regions.

**Extended Data Fig 2.**
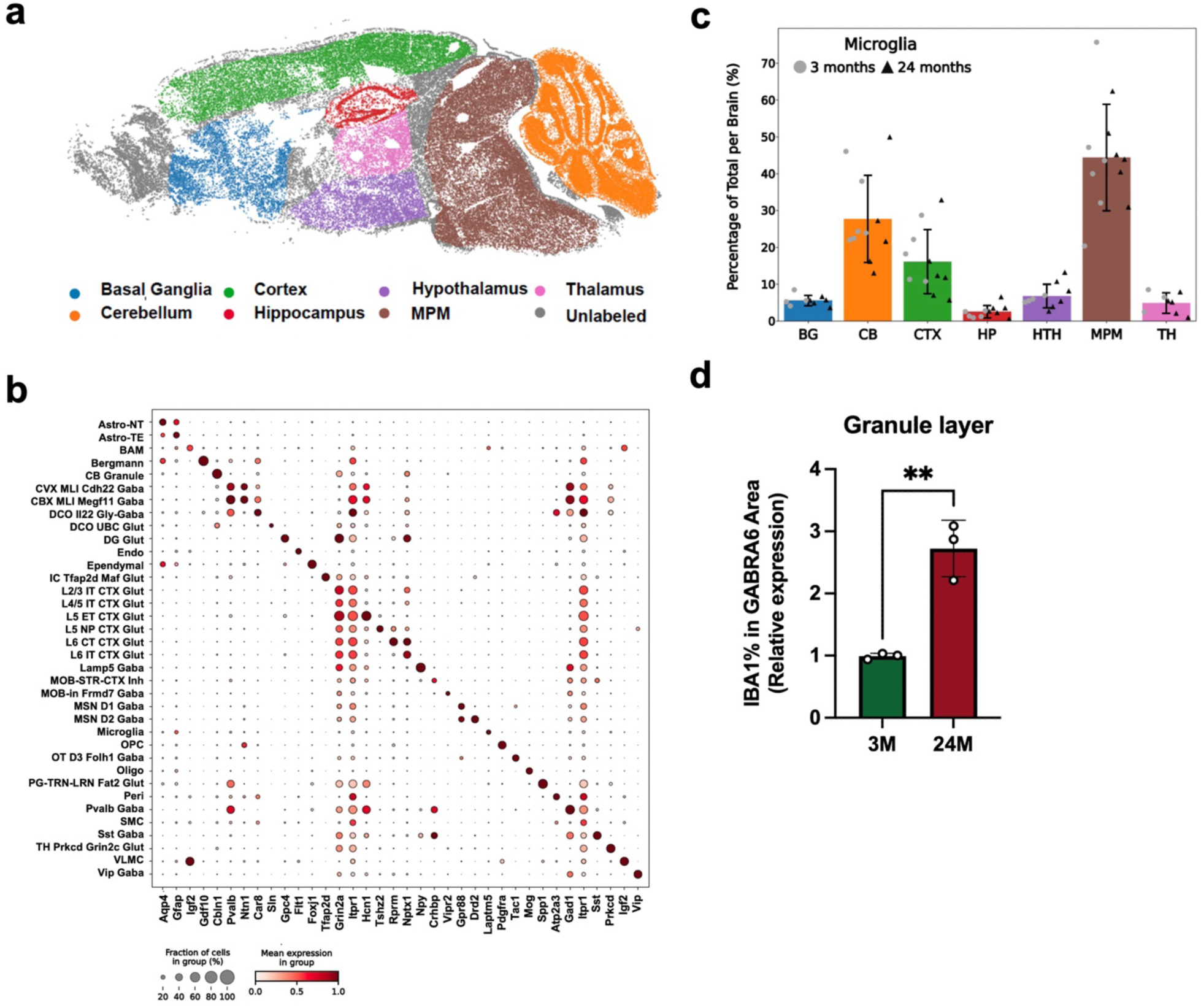
Microglial distribution and markers across different brain regions. **(a),** MERFISH imaging of seven different brain regions, including basal ganglia (BG), cerebellum (CB), cortex (CTX), hippocampus (HP), hypothalamus (HTH), thalamus (TH), and midbrain, pons, medulla oblongata (MPM), with distinct color coding. **(b),** Dot plot showing marker genes for cell subtypes identified in MERFISH, transferred using the Allen Brain Cell Atlas. **(c),** Percentage of microglia in different regions of the brain regions in 3-month-old and 24-month-old mice. **(d)**, Bar plot showing the percentage of IBA1-positive cells (microglia) in the granule layer of the cerebellum (GABRA6-positive area) in 3-month-old and 24-month-old mice (** *p*<0.01).

**Extended Data Fig 3.**
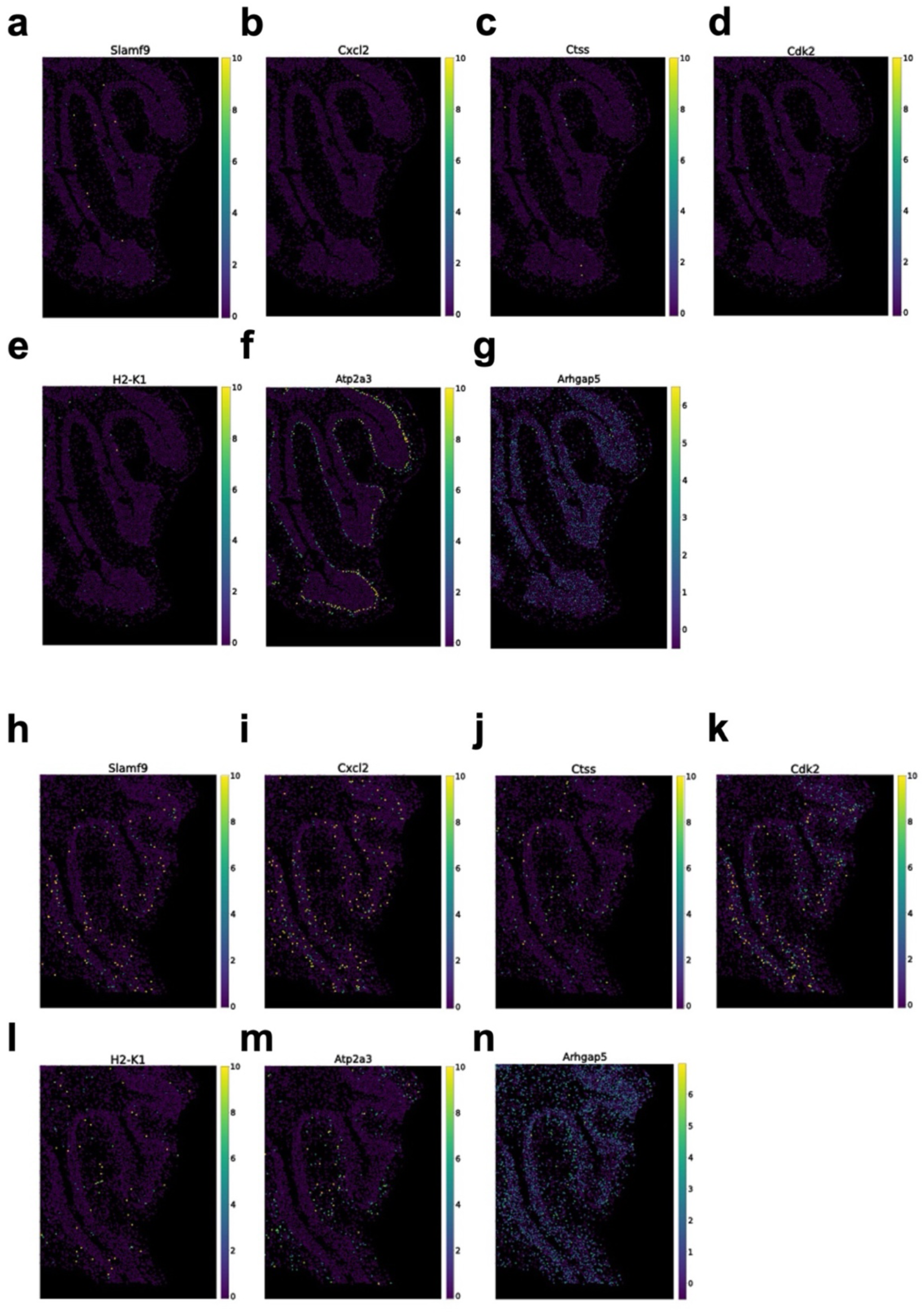
Spatial expression of genes contributing to the neuron-associated microglia score in the cerebellum. **(a-n),** Representative MERFISH images showing spatial expression of *Slamf9* (**a, h**), *Cxcl2* (**b, i**), *Ctss* (**c, j**), *Cdk2* (**d, k**), *H2-K1* (**e, l**), *Atp2a3* (**f, m**) and *Arhgap5* **(g, n)** in the cerebellum of young mice **(a-g)** and aged mice **(h-n)**.

**Extended Data Fig 4.**
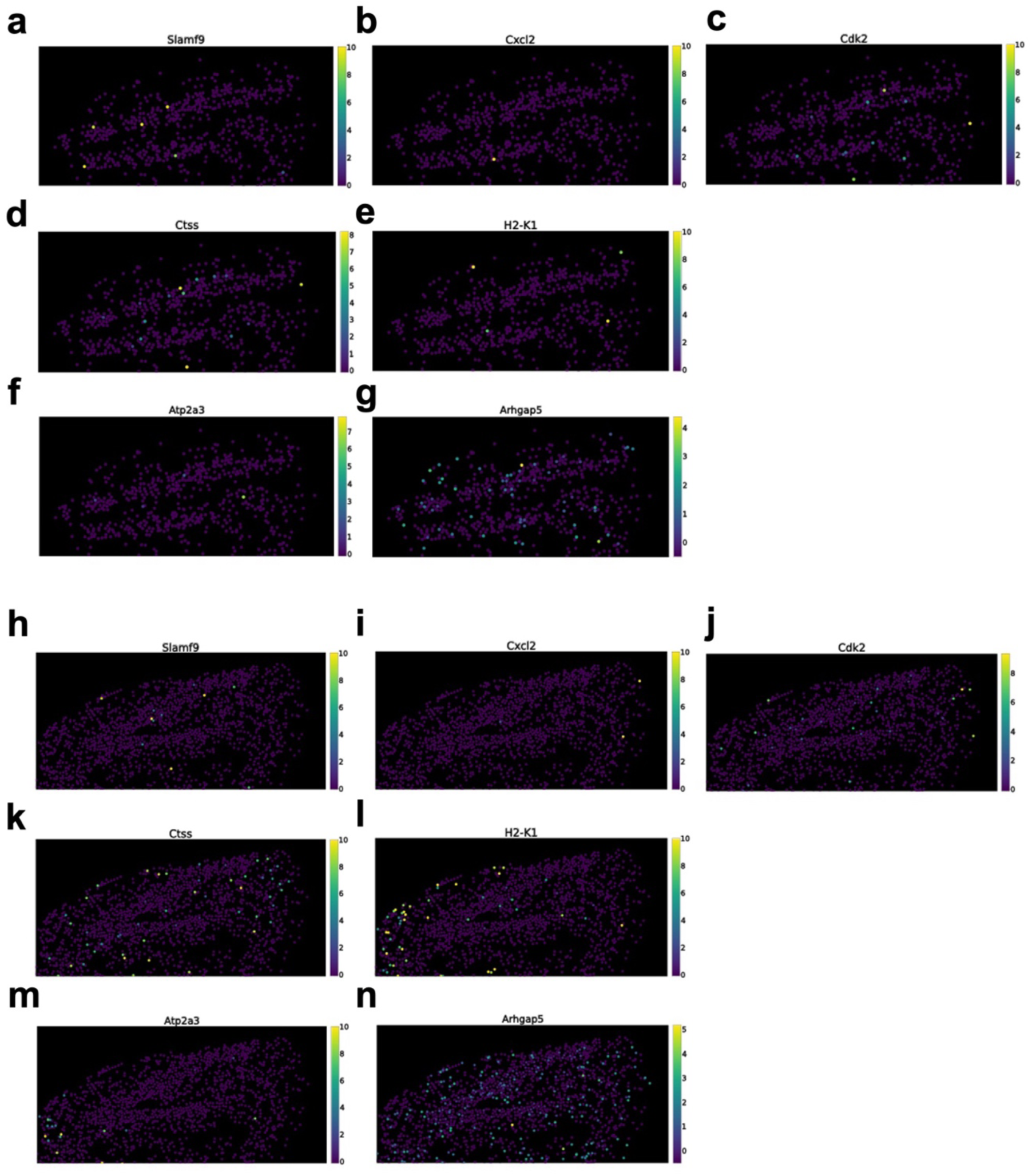
Spatial expression of genes contributing to the neuron-associated microglia score in the hippocampus. **(a-n),** Representative MERFISH images showing the spatial expression of *Slamf9* (**a, h**), *Cxcl2* (**b, i**), *Ctss* (**c, j**), *Cdk2* (**d, k**), *H2-K1* (**e, l**), *Atp2a3* (**f, m**) and *Arhgap5* **(g, n)** in the hippocampus of young mice **(a-g)** and aged mice **(h-n)**

**Extended Data Fig 5.**
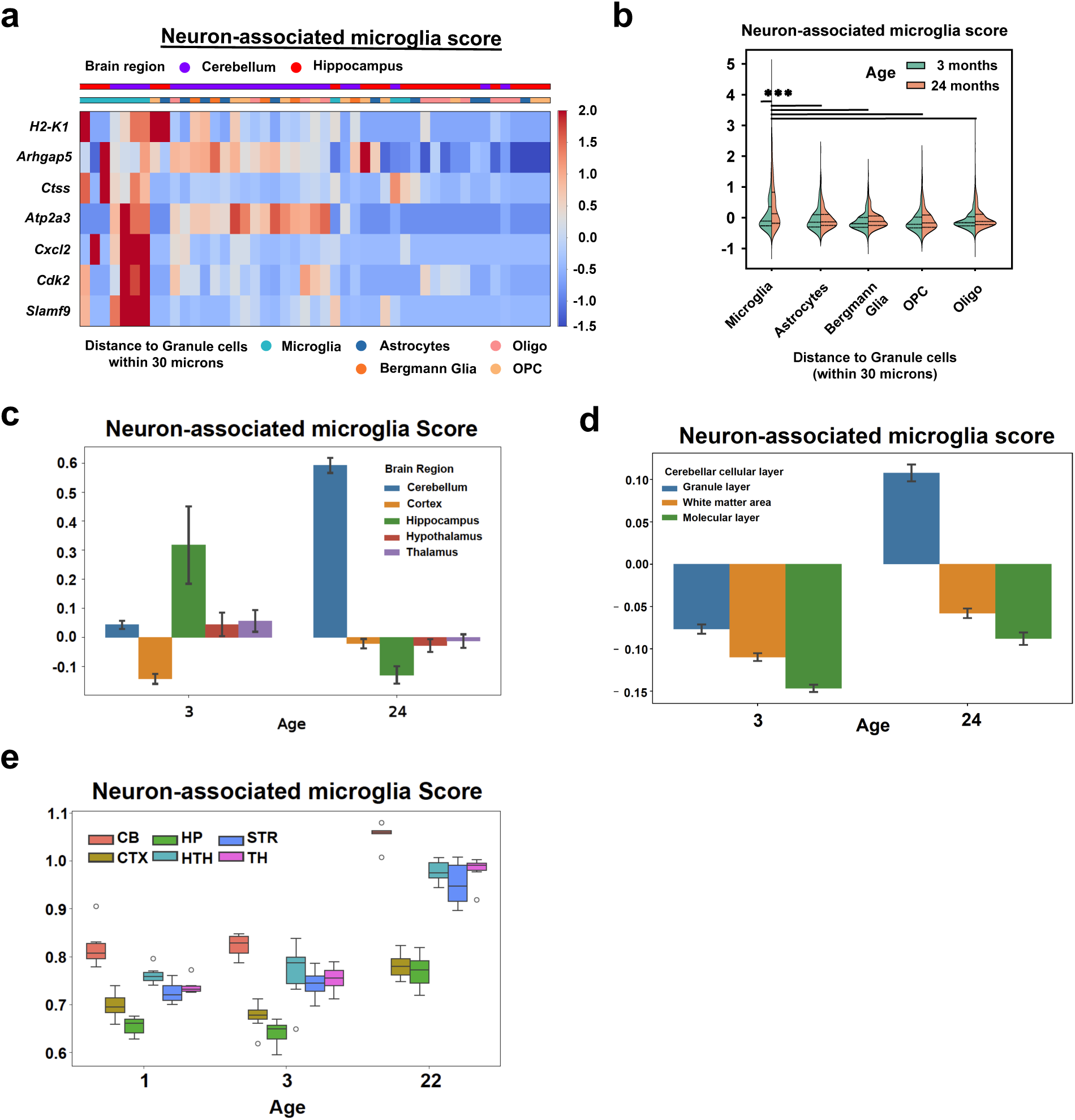
Neuron-associated microglia score across age, brain regions, cell types, and cerebellar layers. **(a),** Heatmap showing the expression of neuron-associated microglia score genes in glial cells within 30 microns of granule cells in the aged cerebellum (purple) and hippocampus (red). **(b),** Violin plots showing significantly higher neuron-associated microglia scores in aged cerebellar microglia compared to young microglia and other glial cells (*** *p*<0.005). **(c-d),** Bar plot showing the neuron-associated microglia score derived from MERFISH images in different brain regions (**c**) and cerebellar cellular layers (**d**) in young and old mice. **(e),** Box plot showing neuron-associated microglial scores from microglial bulk-RNA seq analysis across three age groups (1-, 3-, and 22-month-old) and six brain regions

## Notes

### Competing Interest Statement

The authors have declared no competing interest.

